# Quantification of the diversity sampling bias resulting from rice root bacterial isolation on popular and nitrogen-free culture media, using 16S amplicon barcoding

**DOI:** 10.1101/2022.12.01.518680

**Authors:** Moussa Sondo, Issa Wonni, Agnieszka Klonowska, Kadidia Koïta, Lionel Moulin

## Abstract

Culturing bacteria from plant material is well known to introduce a strong bias compared to the real diversity present in the original samples. This bias is related to cultivability of bacteria, the chemical composition of media and culture conditions. The bias of recovery is often observed but was never quantified on different media using an amplicon barcoding approach comparing plant microbiota DNA extractions versus DNA extracted from serial dilutions of the same plant tissues grown on bacterial culture media. In this study, we i) quantified the culturing diversity bias using 16S amplicon barcode sequencing by comparing a culture-dependent approach (CDA) on rice roots on four popular bacterial media (Tryptone Soybean Agar-TSA-at two concentrations, 10% and 50%; a plant-based media with rice flour; Norris Glucose Nitrogen Free Medium-NGN; and Nitrogen Free -NFb) versus a culture-independent approach (CIA) assessed from DNA extracted directly on root and rhizosphere samples; ii) assessed enriched and missing taxa detected on the different media; iii) use biostatistics functional predictions to predict which metabolic profiles are enriched in the CDA and CIA. A comparative analysis of the two approaches revealed that among the 22 phyla present in the microbiota of the studied rice root samples, only five were present on the culture media approach (*Proteobacteria, Firmicutes, Bacteroidetes, Actinobacteria, Verrucomicrobia*). The *Proteobacteria* phylum was the most abundant in all cultured media samples, showing a high enrichment of gamma-Proteobacteria. The diversity of the combined culture media represented about 1/3 of the diversity of the total microbiota, and its genus diversity and frequency was documented. The functional prediction tool (PiCrust2) detected an enrichment of nitrogenase enzyme in bacterial taxa sampled from Nitrogen-free media, validating its predictive capacity. Further functional predictions also showed that the CDA missed mostly anaerobic, methylotrophic, methanotrophic and photosynthetic bacteria compared to the culture independent approach, delivering valuable insights to design ad-hoc culture media and conditions to increase cultivability of the rice-associated microbiota.

## Introduction

Plants interact continuously with a microbiota that plays an important role in their health, fitness and productivity. In the last ten years, the accessibility to scientist of next generation sequencing (amplicon-based sequencing and metagenomics) at low cost has allowed to describe extensively the diversity of this microbiota on many model and non-model plants (for example in *Arabidopsis* (1) or wheat (2). For rice, many studies, have described its microbiome in different countries and culture practices (3–6). Such wealth of data gives today a good overview of the main bacterial and fungal taxa inhabiting plant tissues underground (roots and rhizosphere) as well as in their aerial parts (phyllosphere and endosphere). The diversity based on amplicon-barcode approaches is mainly based on fragments of ribosomal taxonomic markers as 16S and 18S rRNAs, with a taxonomic resolution often restricted at the genus level. To access and obtain more representativeness of microbial diversity and structure, several studies developed a combination of markers at different resolution levels, from general (16S V3-V4 or V4 for prokaryotes, 18S V4 for microeukaryotes) to more resolutive markers (fragments of *gyrB* or *rpoB* for bacteria, ITS1/ITS2 for fungi) (7–9). The bioinformatic analysis of amplicon barcode data has also encompassed several evolutions, from OTU clustering at different % of identity to more advanced clustering methods using swarming algorithms (10,11), to methodologies inferring the true amplicon sequence variants (12).

Harnessing the diversity of the plant microbiota for example for plant nutrition or tolerance to pathogens relies on the isolation and culturing of the taxonomic and/or functional diversity of the microbiota (13). The capacity to culture and stores such diversity allows to design synthetic communities and test their various compositions on plant growth and health (14,15). Concerning culturomics, different approaches have been developed to capture the bacterial diversity of plant microbiota, including for example culture media supplementation with various compounds, simulated natural environments, diffusion chambers, soil substrate membrane systems, isolation chips, single cell microfluidics (reviewed in (16)), or using limiting dilutions in plates coupled with dual barcodes processing (17). Significant improvements in diversity sampling have also been achieved by popular media supplementation with plant compounds or plant-based media, and microbiologists continue to develop alternative culture methods in order to capture rare and unculturable plant-associated microorganisms (16).

Several functional prediction tools have recently been developed to predict the enrichment of functions in metagenomes and even 16S amplicon barcoding data (as PiCrust2 (18)). In theory, such tools could allow to identify which metabolic functions (and ecology) are enriched in culture independent approaches compared to culturable ones, in order to better define culture media or culturing conditions to trap them.

It is well accepted in the scientific community of microbiologist that popular non-selective bacterial media (as Lubria Broth -LB, R2A, Nutrient Agar, Tryptic Soy Agar - TSA) introduce a strong bias in the recovered sampled diversity from plant tissues, but this bias is to our knowledge always observed but never quantified and documented in terms of proportions, by using NGS amplicon-based technologies. Other media have been designed to isolate dinitrogen-fixing bacteria (Norris-Glucose Nitrogen Free medium-NGN, Nitrogen Free-NFb medium, (19) with success, but again without knowing which proportion of the nitrogen fixing community was captured compared to the real diversity of nitrogen fixers.

In this study we used 16S amplicon barcode sequencing on DNA extracted directly from roots and rhizosphere of rice (culture-independent approach-CIA) as well as on DNA extracted from mass bacterial culture of several dilutions of the same plant samples plated on a popular media for plant-associated bacteria (TSA at 10 and 50%), a plant-based media (rice flour) and two nitrogen-free media (NFb, NGN), in order to i) quantify the bias of diversity from culturing on popular and nitrogen-free media versus real bacterial diversity, using last bioinformatic methodologies to identify the bacterial diversity (Amplicon-Sequence Variant - ASV and OTU assessment by swarming methodology) ; ii) determine proportions of enriched bacterial genera per medium, iii) use functional predicting tools on amplicons data to identify specific metabolic functions or bacterial capacities that are present in the rice root microbiota but missing from our culturable approach. Our hypothesis was that using amplicon-sequencing on DNA extracted from bacterial growth after plating serial dilutions of plant tissues would decrease the problem of loss of slow-growing bacteria compared to fast-growing ones, due to the deep-sequencing approach, and would give a much better assessment of culturable bacterial diversity and percentage of recovery compared to the real diversity.

## MATERIAL & METHODS

### Rice roots sampling and processing

*Oryza sativa* ssp *indica* cv FKR64 plant roots were collected in a rice field near Bama village (West of Burkina Faso, Kou Valley, 10.64384 North, -4.8302 East). This field was already studied in a previous study and is described in (6). Rice plants were sampled at the panicle initiation growth stage, with three sampling points chosen 10 m apart from each other, where roots were collected from three plants (20 cm apart). Roots were hand-shaken to remove non-adherent soil. Ten roots per plant from the same sampling point were pooled to obtain three final samples in 50 ml Falcon tube containing 30 ml of sterile PBS buffer, and vortexed 5 min to separate the rhizospheric soil from the roots. Roots were removed with sterile forceps and placed in new 50 ml Falcon tubes. From this treatment step, the rhizosphere (Rh) and roots (Rt) samples were manipulated separately (see Figure 1). The rhizosphere soil in PBS was vortexed for 10 sec and then two samples of 1 ml of the rhizosphere suspension were taken after 15 sec and placed in two separate 2 ml Eppendorf tubes to be used for either bacterial cultivation-dependent (CDA) or cultivation independent approaches (CIA) for diversity estimation by direct 16S amplicon barcoding approach. Similarly, washed roots, were cut into 2 cm fragments, and separated into two 2 ml Eppendorf tubes, for CDA and CIA approaches.

**Figure 1.**
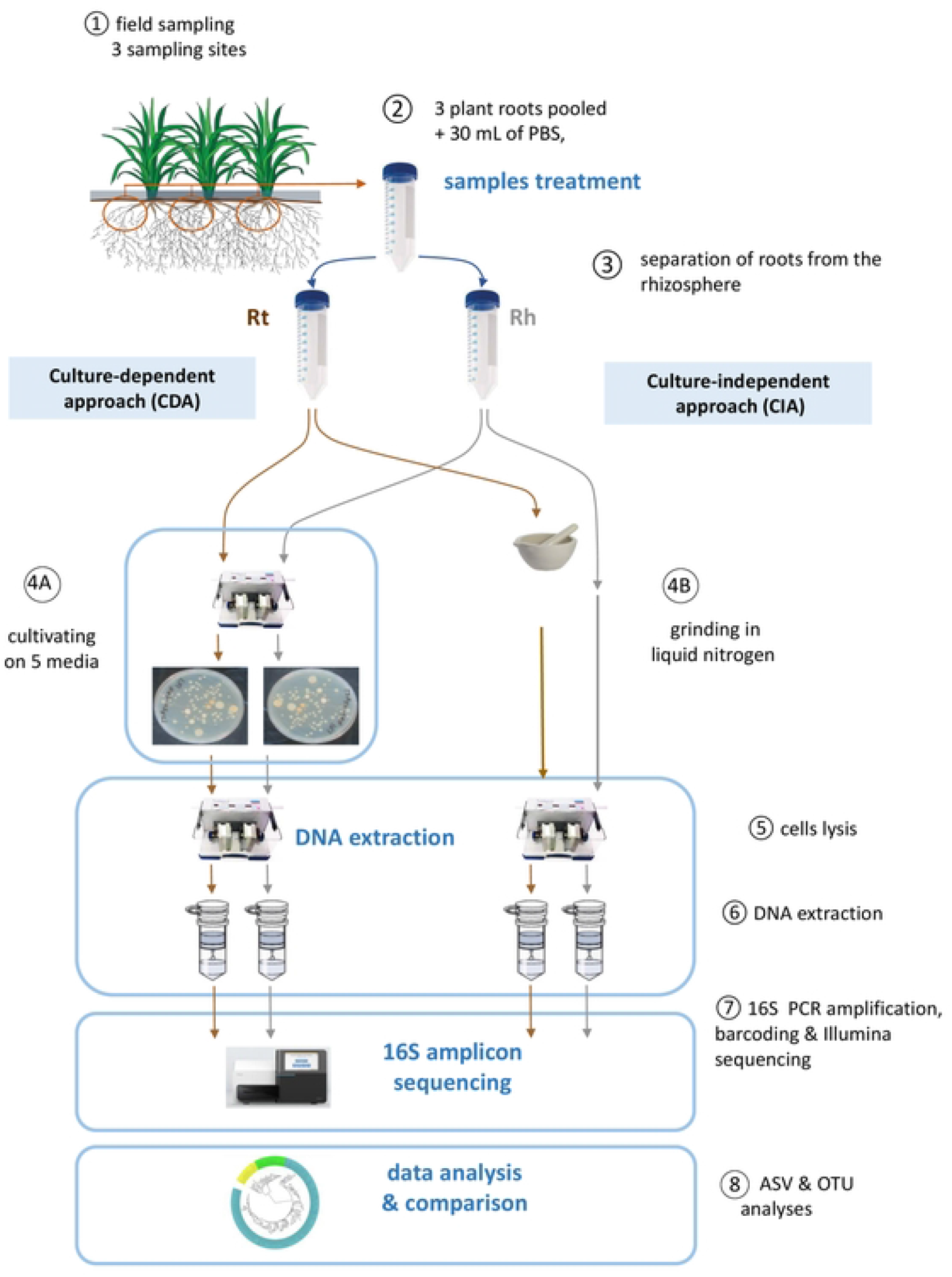
Schematic representation of the protocol of sampling and processing for 16S amplicons libraries.

### Bacterial culture isolation media

Four culture media employing different carbon and nitrogen sources were used to maximize the diversity of isolated bacteria. First, the non-selective Tryptic Soy Agar (TSA, Sigma) medium was used at two concentrations 10% (TSA10) and 50% (TSA50). It contains digests of casein and soybean meal, NaCl and agar. In addition, two nitrogen-free media were used for the isolation of potential nitrogen fixers, semi-solid NFb (19) and Norris Glucose Nitrogen Free Medium (NGN, M712, (20)). NFb was used as semi-solid medium, which allows the development and growth of free nitrogen-fixing bacteria, due to their growth at an optimal distance for micro-aerobic condition favourable for nitrogen fixation (19). Finally, we included a plant-based medium, rice flour (RF) usually used for isolation of fungal rice pathogens (21). The composition of above culture media was as follows: NGN (g/L), 1.0 K_2_HPO_4_, 1.0 CaCO_3_, 0.2 NaCl, 0.20 MgSO_4_·7H_2_O, 0.01 FeSO_4_·7H_2_O O, 0.005 Na_2_MoO_4_·2H2O, and a carbon source was glucose (10 g/L) and pH 7; NFb : (g/L), 0.5 K2HPO4, 0.2 MgSO4.7H2O, 0.1 NaCl, 0.02 CaCl2. 2H2O, 4.5 KOH, 5 Malic acid, 2 mL of micronutrient solution ((g/L) 0.04 CuSO4.5H2O, 0.12 ZnSO4.7H2O, 1.40 H3BO3, 1.0 Na2MoO4.2H2O, 1.175 MnSO4. H2O), 2 mL of bromothymol blue (5 g/L in 0.2 N KOH), 4 mL of FeEDTA (solution 16.4 g/L), 1 mL of vitamin solution ((mg/0.1 L) 10 biotin, 20 pyridoxal-HCl) and pH adjusted to 6.5; RF (g/L): 20 rice flour (prepared seeds of FKR64 rice variety), 2.5 Yeast Extract. Solid and the semi-solid media were obtained by adding 2% and 0.16% g of agar, respectively.

### Cultivation-dependent (CDA) & and independent (CIA) approaches

For the CDA, roots (200 mg) and rhizosphere (200 mg) were transferred into PowerBead Tubes from the DNeasy PowerSoil kit (QIAGEN) where 1 mL of PBS buffer was added, and grounded in a TissueLyser II (QIAGEN) for 2 min (Figure 1). Dilutions (10^−2^ to 10^−5^) were performed and 50 µL of each dilution were spread on solid culture media (TSA 10%, TSA 50%, NGN, RF). For NFb medium, 50 µL of the 10^−1^ root and rhizosphere suspensions were inoculated in 20 mL tubes containing 10 mL of NFb semi-solid medium. Each dilution was inoculated (on plates or in tubes) with 4 replicates. After 2 to 5 days of incubation (depending on the culture medium) at 28°C, plates were examined and dilutions selected for further processing (details in see Supplementary Table S1). For selected dilutions, cultivable bacteria were recovered from Petri plates by adding 1 ml of sterile distilled water, scraping and mixing bacterial colonies. Obtained bacterial suspension from the same dilution plates, were collected with a pipette and transferred to sterile 15 ml Falcon tubes. For the NFb medium, the bacteria which have grown in a form of a ring at 0.2-0.3 cm distance below the surface of the medium were collected. Bacterial suspensions were stored at -20°C until DNA extraction. The number of cultivable bacteria in obtained suspensions was roughly estimated by measuring the optical density (OD) at 600 nm for all suspensions and adjusted to 10^6^ (assuming that OD600 nm of 1 corresponds to 1×10^8^ bacteria per mL). The volumes collected from the samples were centrifuged 10 min at 14 000 rpm, and the pellets obtained were used for DNA extractions.

For the cultivation-independent approach (CIA), pooled roots were homogenized in liquid nitrogen using a mortar with pestle, while the pooled rhizosphere samples were used directly for DNA extractions (Figure 1). A mass of 250 mg was used for both sample types for DNA extractions.

### DNA extractions

Cultivable bacteria suspensions (roughly 10^6^ cells) and ground roots and rhizosphere (250 mg) were transferred to PowerBead tubes (DNeasy PowerSoil, Qiagen) containing C1 buffer and homogeneised in a TissueLyser II (Qiagen) at 240 rpm for 2 × 1 min. The extraction was then performed following the protocol provided by supplier.

### 16S amplicon-barcoding data production

Quality control of DNA, PCR amplification, library construction and and MiSeq Illumina sequencing were performed by Macrogen (Seoul, South Korea) using bacterial primers 337F (16S_337F, 5’-GACTCCTACGGGAGGCWGCAG-3’) and 805R (16S_805R, 5’-GACTACCAGGGTATCTAATC-3’) to amplify the V3-V4 region of the 16S rDNA gene (22). The sequencing data (fastq) for this study have been made accessible in the ENA (https://www.ebi.ac.uk/ena) database under the Bioproject (study accession number) PRJEB55863 (ERP140807).

### Bioinformatic analysis of 16S amplicons

For this article we performed all diversity analyses using an Amplicon sequence variant (ASV) detection approach (DADA2 pipeline), but we also performed a comparison of diversity with a OTU clustering method (based on FROGs, (23)).

For ASV analysis, raw amplicon barcoding data were demultiplexed and processed using the Bioconductor Workflow for Microbiome Data analysis (24). This workflow is based on DADA2 (12) that infers amplicon sequence variants (ASV) from raw sequence reads. Forward and reverse reads were trimmed at 20 bp, respectively, to remove primers and adapters, and quality-truncated at 280 and 205 bp respectively. The function dada2 denoise-paired with default parameters was used to correct sequencing errors and infer exact amplicon sequence variants (ASVs). Then forward and reverse corrected reads were merged with a minimum overlap of 20 bp, and the *removeBimeraDenovo* from DADA2 was used to remove chimeric sequences. Eighty two percent of reads passed the chimeric filter. Numbers of reads filtered, merged and non-chimeric are indicated in supplementary Table S2. A mean of 58.6% of reads passed all filters (denoising, merging, non-chimeric), with a minimum of 15347 and a maximum of 31134 reads in filtered libraries.

ASV were then assigned at taxonomic level using the DADA2 *AssignTaxonomy* function, using the Silva 16S reference database (silva_nr_v132_train_set). We subsequently filtered out plasts (especially mitochondria from root samples) to keep only SVs assigned to the Bacteria or Archaea kingdoms. A last filtering was made to remove ASV with less than 10 reads occurrence across all libraries. A dataset of 1677 ASV was used for subsequent diversity analyses. A neighbor-joining phylogenetic tree of the 1677 ASV was constructed using MEGA11 (25), by first aligning ASV sequences with MUSCLE (26) and then building a neighbour joining-tree based on a distance matrix corrected with Kimura 2P method. Metadata and ASV tables and the phylogenetic tree were uploaded to NAMCO server for downstream microbiota diversity analyses (https://exbio.wzw.tum.de/namco/, (27). NAMCO is a microbiome explorer server based on a set of R packages including Phyloseq for diversity analyses (28) and PICRUST2 for functional predictions (18). Circular phylogenetic tree annotations and mapping were produce with iTOL (29). Additional R scripts for the DADA2 pipeline, Phyloseq, and the production of figures are freely available on Github (https://github.com/lmoulin34/Article_Moussa_culturingbias).

For the OTU clustering approach, the FROGs pipeline ((23); http://frogs.toulouse.inra.fr/) was used in Galaxy environment. After demultiplexing and pre-processing, reads were clustered into OTU using the swarming method with default parameters, then chimeric sequences were removed and OTU were affiliated to taxonomic levels using the same Assign taxonomy tool as described above.

## RESULTS

### Quality filtering & diversity indices of 16S amplicon libraries (CIA versus CDA)

We first assessed the quantity and quality of reads produced for each amplicon library originating from direct rice root or rhizosphere genomic DNA extraction (CIA) or from DNA extracted from grown cultures (CDA) of the same samples on bacterial culture media. A range of 24000 to 44000 reads (mean at 36120) was obtained for all 16S amplicon libraries (Supplementary Table S2). Rarefaction curves (Figure 2A) showed saturation of sampled diversity for each library, with a clear difference between the CIA microbiota reads (much higher in alpha diversity) compared to CDA.

**Figure 2.**
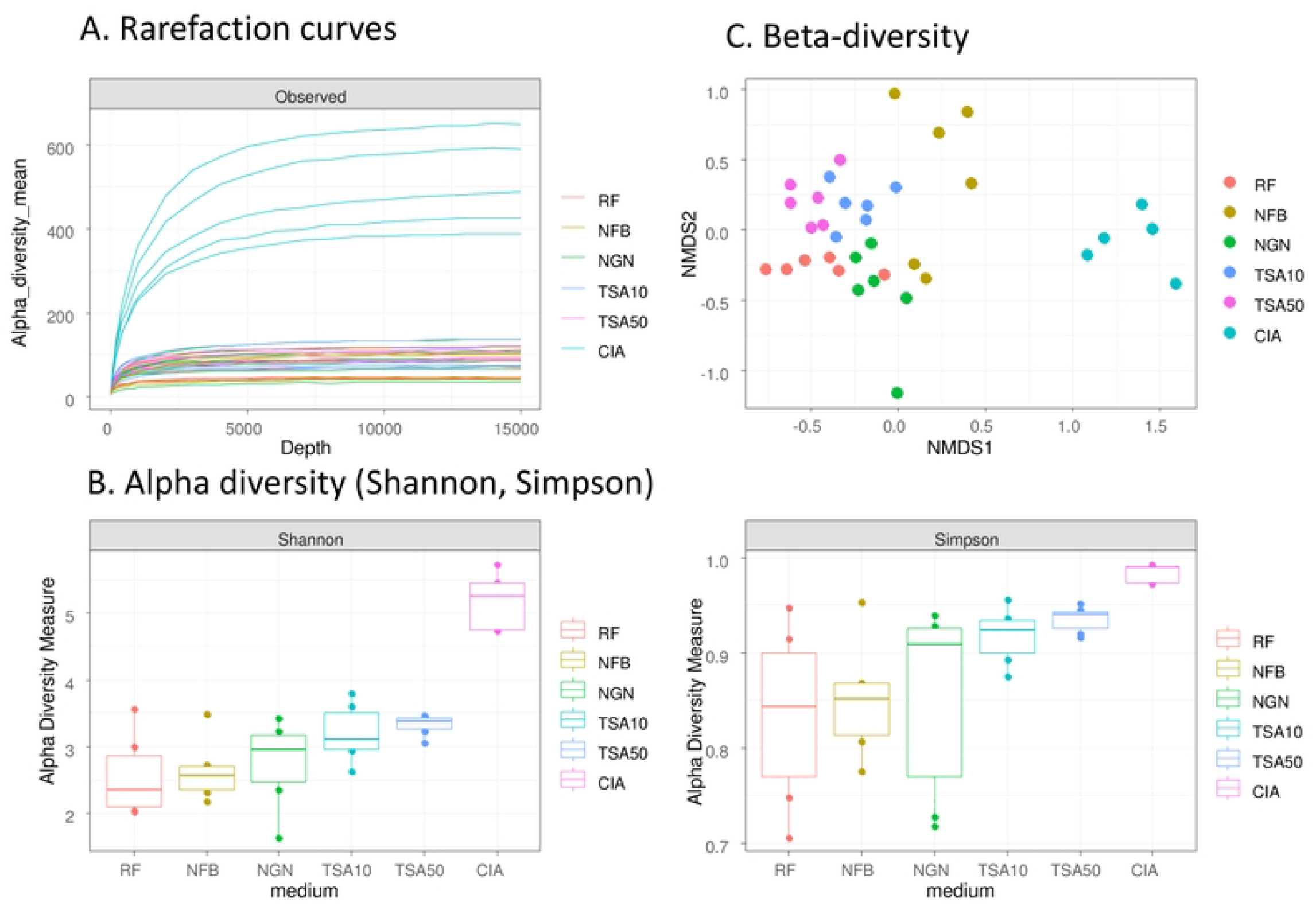
Rarefaction curves (A), alpha (B) and beta diversity (C) of 16S amplicon libraries.

After DADA2 pipeline processing, we obtained 2712 amplicon sequence variants (ASV) that were assigned at taxonomic level using the Silva database. One library (S36) was removed from the analysis (from CIA) as it showed only 3 ASV. For remaining libraries, ASV were filtered on their abundance (cumulated reads ≥ 10 among all libraries) and mitochondria, chloroplast and eukaryote reads were removed (remaining ASV=1646).

We first compared the diversity obtained from root and rhizosphere samples. As expected, the richness was more important in the rhizosphere microbiota (Rh) compared to the root one (Ro) (Supplementary Figure 1). Beta-diversity of the same samples did not show significant differences among Ro and Rh libraries (Supplementary Figure 1). Such results can be explained by the fact that we did not surface disinfect nor removed the rhizoplane from roots, so the rhizosphere (adherent soil to root) and the root (rhizoplane + endosphere) from the same samples did not show high differences. As the focus of this study was on the comparison between the plant microbiota and the diversity obtained from a culturable approach on different media, we pooled Rh and Ro data from the same plant samples for all subsequent analyses.

Analysis of the alpha diversity of all sequences in the culture media used (TSA10, TSA50, NGN, NFb, RF) compared to the CIA microbiota sequences showed that the microbiota had a higher specific richness compared to the diversity sampled on each media (Figure 2B). The TSA, RF and Nitrogen-free media generated nearly identical richness (Figure 2B). The richness sampled from each medium represents about 25% of the diversity of the microbiota (TSA10: 22.45%, TSA50: 24.45%, NFb: 20.41%, NGN: 22.45%), except for RF (14.28%) which captured less diversity. NMDS on beta diversity analyses showed no overlap between ASV obtained from the different media and the microbiota (Figure 2C), while a large overlap was observed for TSA10 and TSA50, which was expected since it is the same medium used at two different concentrations. Beta diversity from media as NGN and NFb showed poor overlap with other culture media.

### Culturable sampled diversity: comparison between ASV and OTU

We also analysed our amplicon barcoding reads using an OTU-clustering approach (FROGs pipeline, using the swarming method to merge reads into OTU). This approach produced 1023 OTU after quality filtering criteria (same as for ASV analysis). We then assessed if the diversity obtained by OTU gave the same % of recovery of diversity compared to ASV. In Table 1 we present the number of ASV and OTU obtained from the culture-dependent approach (CDA) and from the culture-independent approach (CIA), as well as number of classes, orders and families represented in each. The ASV analysis produced more alpha diversity (38% more) than the OTU one. Such higher diversity was observed at different taxonomic levels: class (ASV:50; OTU:38), order (ASV: 124; OTU:67), and families (ASV:219; OTU:119). Given this result we did all subsequent analyses with ASV-analysed data as it performed better at capturing the diversity of our 16S amplicons libraries. In both analyses the shared diversity between CDA and CIA was relatively low (7% for ASV, 22% for OTU). Thus, we recovered from the culturable approach many bacterial taxa that were undetected in the amplicon sequencing performed on gDNA extracted from roots or the rhizosphere, while on the other hand only a small proportion of the root bacteria were able to grow on our culture media.

**Table 1.**
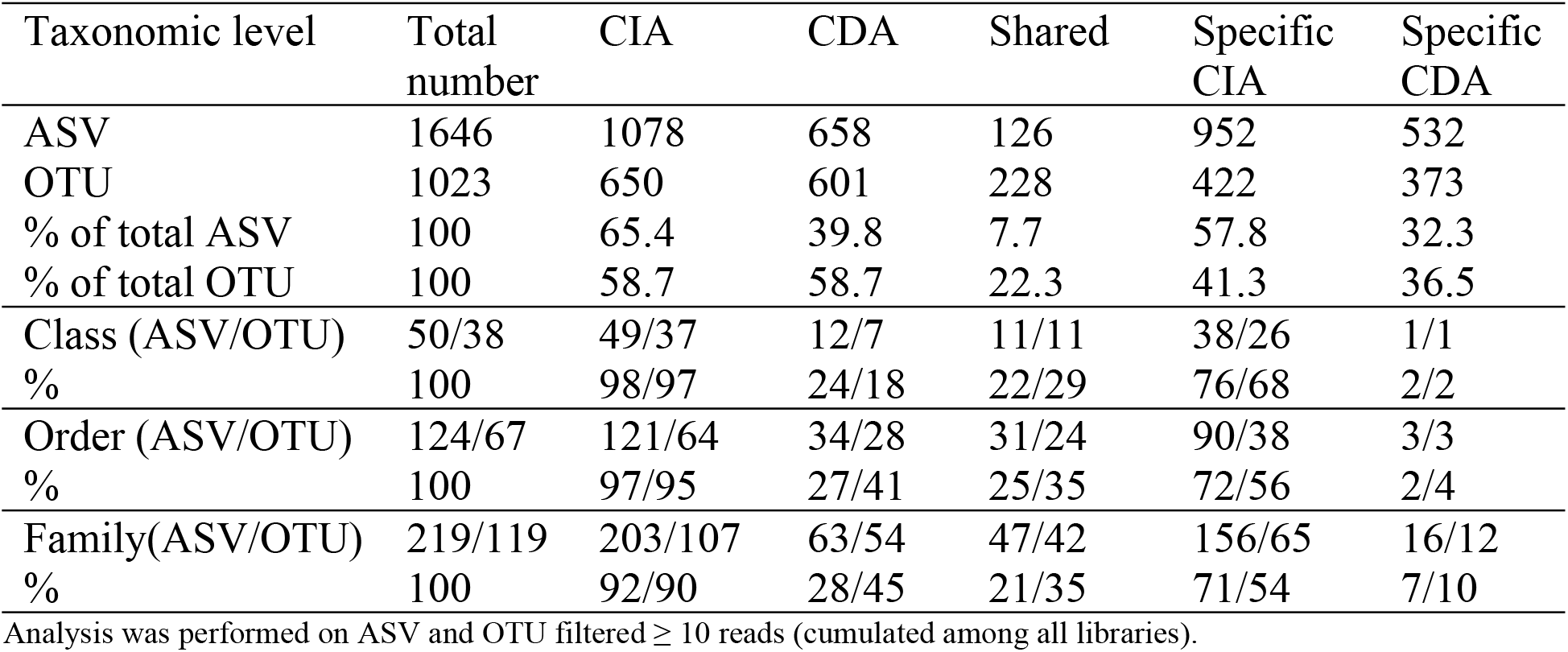
Comparison of diversity in culturable dependent (CDA) and culturable independent (CIA) (CIA) approaches, using ASV or OTU analysis.

### Comparison of taxonomic diversity between microbiota (CIA) and culture media (CDA)

Taxonomic binning was performed at different taxonomic levels for the top 30 Phyla and the top 25 Classes, Orders and Genera (Figure 3A to 3D). Phylum distribution showed a dominance of *Proteobacteria, Bacteroidetes* and *Firmicutes* in all libraries, with a clear higher diversity of phyla for the CIA samples. If we identified 22 phyla in the microbiota of rice root samples, only 5 out of the 22 were present in the CDA (*Proteobacteria, Firmicutes, Bacteroidetes, Actinobacteria, Verrucomicrobia*). The proteobacteria phylum is the most abundant in all the samples with a greater proportion on the rice flour culture medium. At class level, the difference in diversity is even more visible with *Gammaproteobacteria, Alphaproteobacteria* and *Bacteroidia* dominating in the CDA, while in the CIA a large diversity of classes are present (Figure 3B). At the order level, the CIA shows as expected a large diversity while the CDA data are dominated by the *Enterobacteriales, Betaproteobacteriales, Rhizobiales* and *Flavobacteriales*. Finally, on the top 25 genera, differences among CDA libraries appear clearly with the exception of the *Enterobacter* genus which is enriched in all (though to a less extent for NFb) (Figure 3D). In the CIA, the most represented genus is *Devosia*. To better visualize the sampled diversity distribution, we built a phylogenetic tree of ASV (diversity labelled at class level) and mapped their distribution and abundance in the different conditions (coloured outer circles) (Figure 3E). Such representation allows clearly to spot which taxa diversity is sampled and over-represented with the media used in the CDA (for example *Gammaproteobacteria* in blue or *Firmicutes* in pink), and which whole parts of bacterial diversity were missed compared to the CIA (for example *Patescibacteria, Armatimonadetes, Deltaproteobacteria, Planctomycetes, Chloroflexi*).

**Figure 3.**
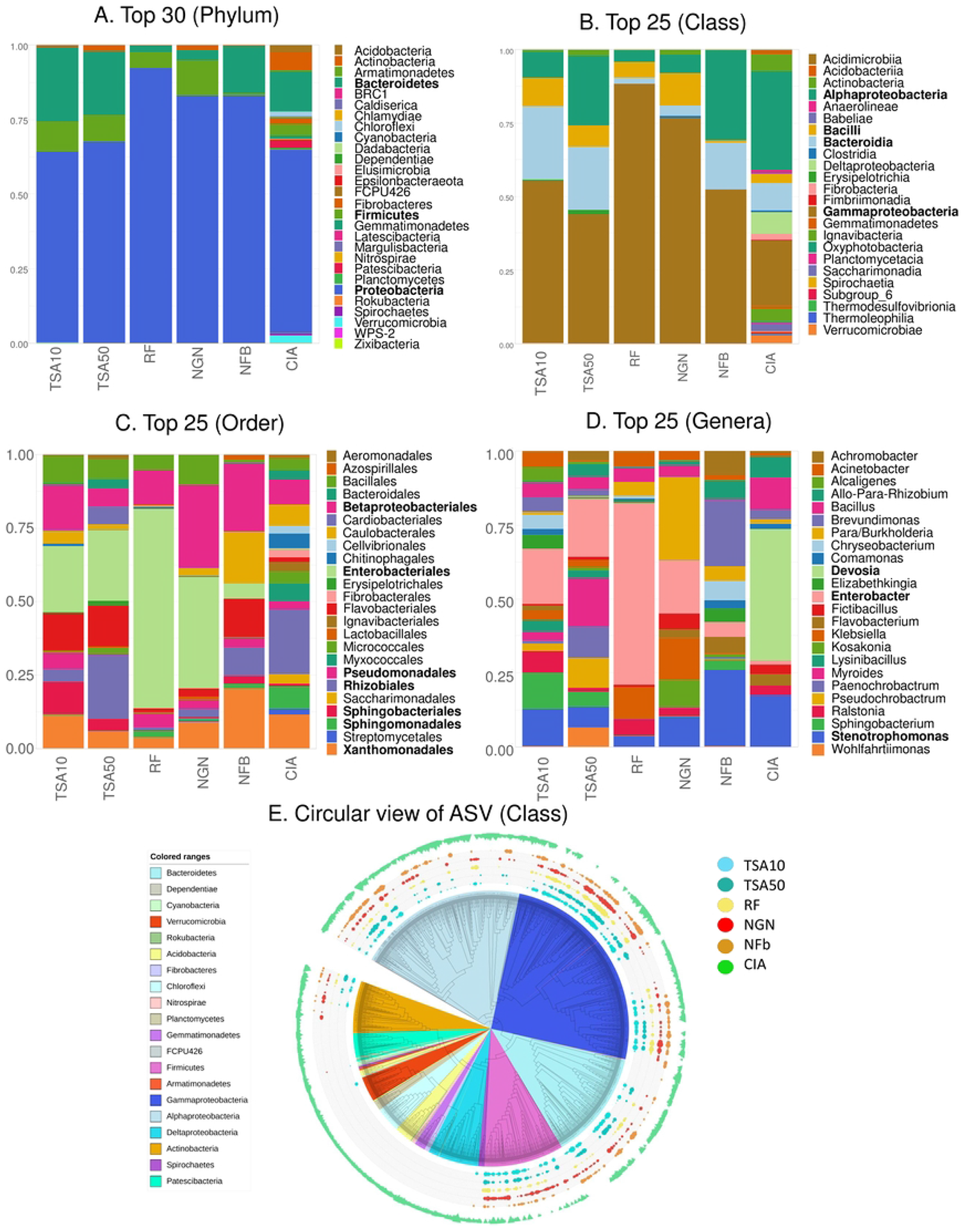
Taxonomic binning of ASV at Phylum (A, top 30), Class (B, top 25), Order (C, top 25) and Genus (D, top 25) level, and circular phylogenetic tree of ASV with class-rank distribution among CIA and CDA (E).

### Statistical differential analyses between CIA and CDA at class and genus level

We performed Kruskal-Wallis test (p cut-off at 5%, with Bonferroni multiple test correction method) to identify classes of bacteria with significant differences among CDA and CIA conditions. The Wilcoxon tests identified 45 classes of bacteria above the significance cut-off level, among which 37 were present only in the CIA (Supplementary Table S3), including in the top 10 most frequent class taxa: *Ignavibacteria, Saccharimonadia, Fibrobacteria* and *Acidobacteriia*. Four classes were present in both the CIA and CDA: *Alphaproteobacteria, Gammaproteobacteria, Bacteroidia* and *Actinobacteria*; the *Alphaproteobacteria* and *Gammaproteobacteria* being the most represented in the CIA and CDA, respectively (also visible on Figure 3B & 3C).

Then we performed differential analyses on the mean relative abundance of bacterial genera in each condition, using a Kruskall-Wallis test at a p-value cut-off of 5%. Table 2 shows the 50 most abundant genera in the CIA microbiota and their mean relative abundance in each media dataset (whole dataset available as Supplementary Table S4). Among 20 most frequent genera in the rice root microbiota (CIA) eleven were detected in the CDA. These were *Devosia* (8.25% of all genera), obtained from TSA10, TSA50 or NFB media; followed by *Pseudoxanthomonas* (3.62%) found in all but RF media condition, then *Stenotrophomonas* (3.36%), *Bacillus* (2.29%), *Pseudomonas* (1.42%) and *Allo/Neo/Para/Rhizobium* (1.3%) found in all media; and finally *Sphingopyxis* (2.1%) detected in TSA50; *Streptomyces* (1.48%) in NGN and *Pseudolabrys* (1.47%) in NFb. We built Venn diagrams on shared and specific diversity at ASV (Figure 4A) and genus level (Figure 4B-C). Among the 244 genera from the CIA, 173 (71%) were absent from the culturable approach, while 71 were shared (29%) and 70 others are CDA-specific (Figure 4B). We also compared the genus diversity sampled by each media of the CDA, and listed specific genera obtained for each media, on the Venn diagram in Figure 4C.

**Table 2.**
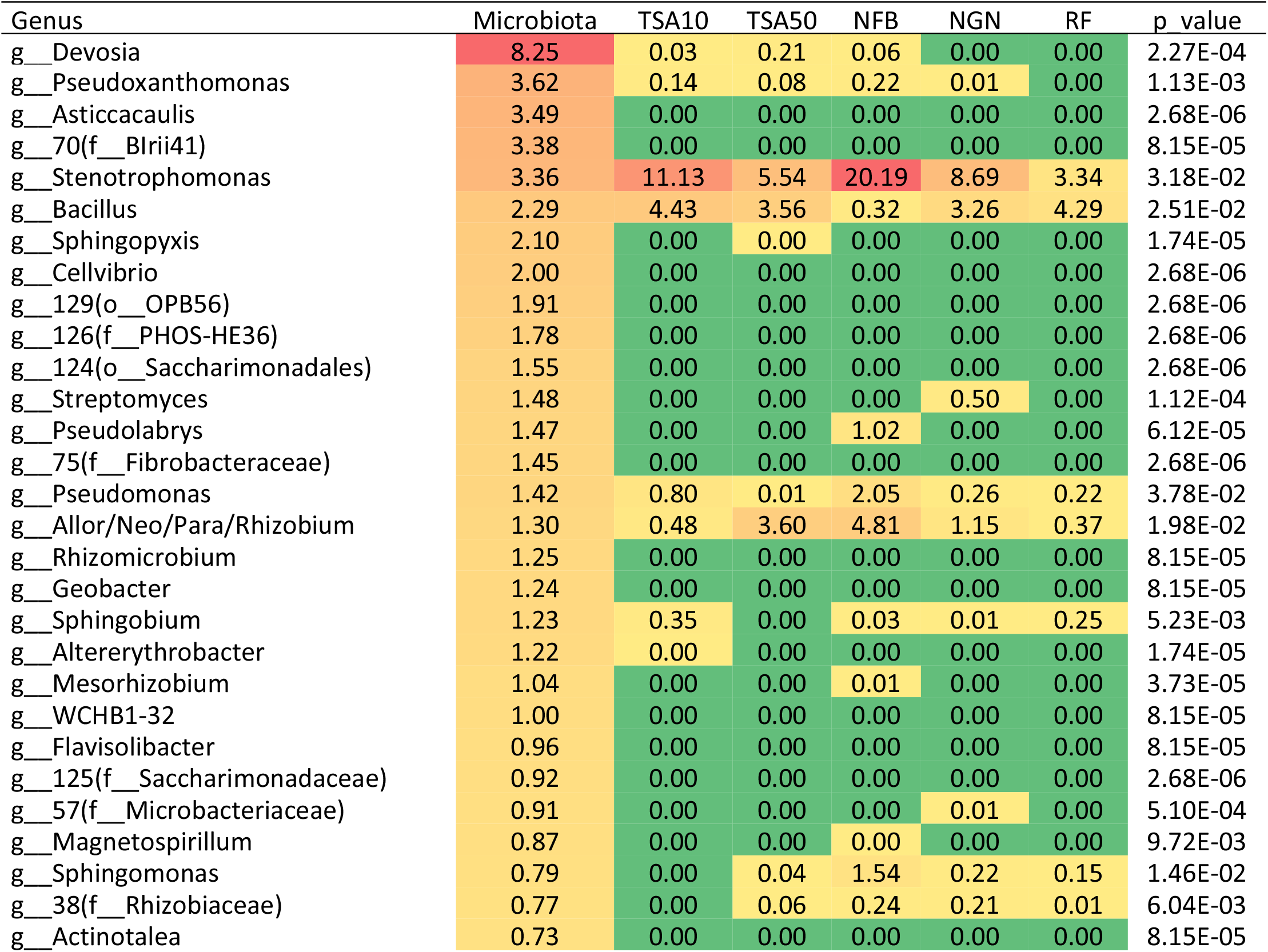
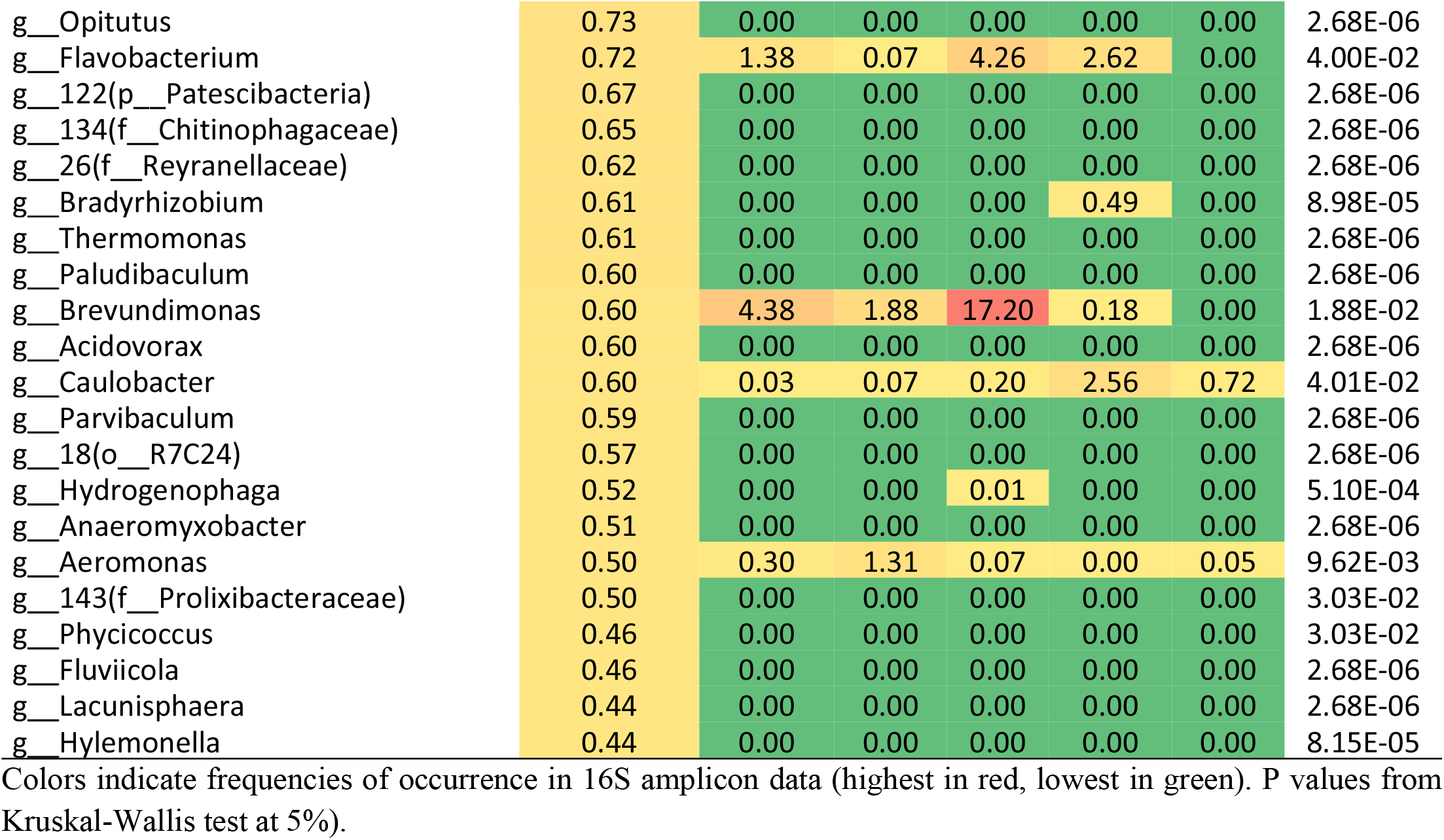
List of top 50 most abundant genera in rice microbiota (CIA), and occurrence in the media of the culturable-dependent approach.

**Figure 4.**
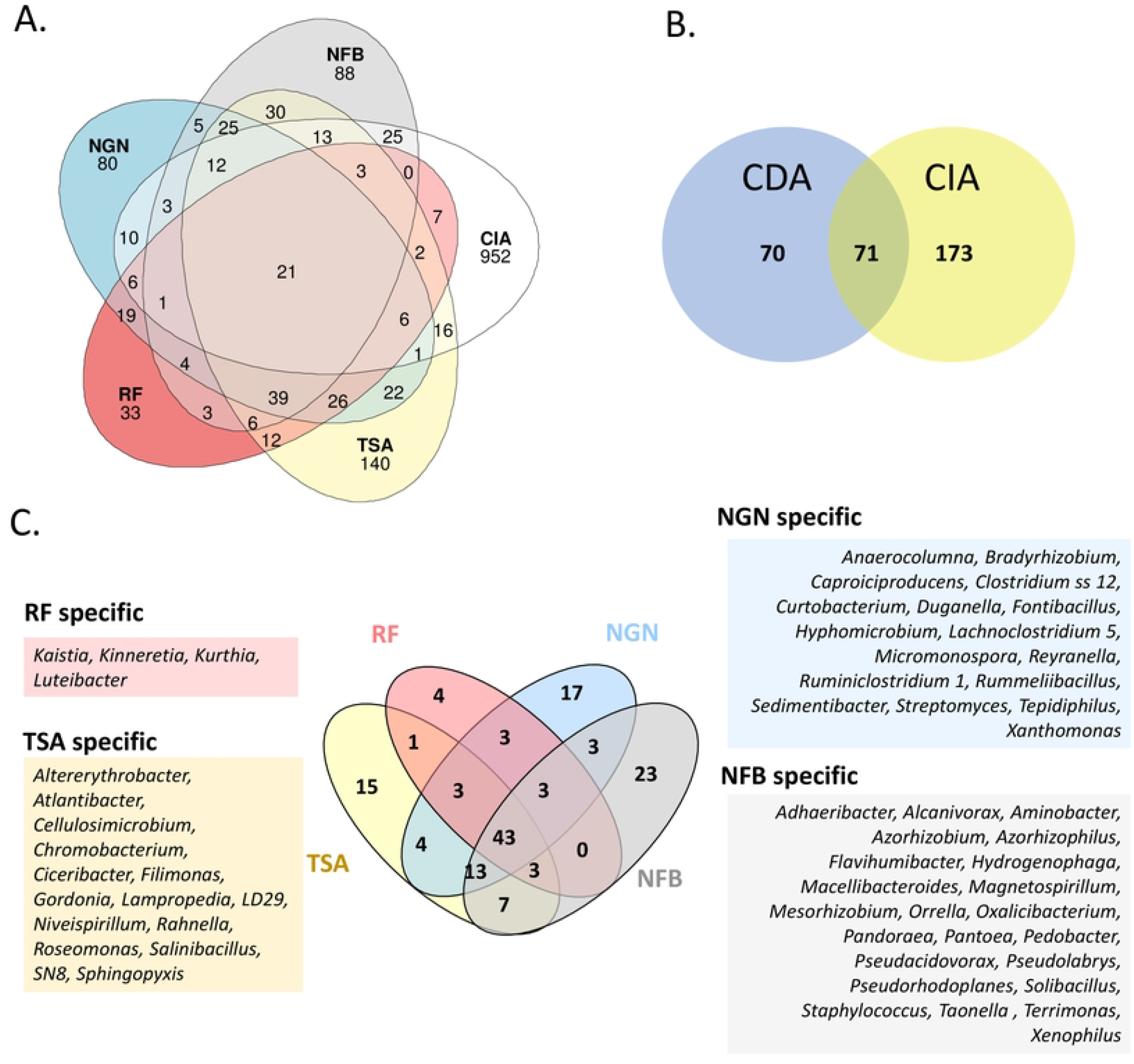
Venn diagrams of diversity between CDA and CIA at ASV level (A), genus level (B), and between culture media used for the CDA (C). Specific genera obtained on a given culture media are listed in (C).

To document which genera are the most frequent in culturable approach for each medium, a table of the 20 most statistically frequent genera (Kruskal-Wallis test at 5%) obtained for each media of the CDA is given in Table 3. In this top 20 most frequent genera, several appeared in all media: *Enterobacter, Stenotrophomonas, Bacillus, Sphingobacterium, Klebsiella, Brevundimonas, Rhizobium*, all known to be fast-growers on rich media and reported in the literature as containing plant-inhabiting species. Nitrogen-free media additionally sampled species known as nitrogen-fixing PGPR: *Azospirillum, Para/Burkholderia, Bradyrhizobium; Sphingomonas*, among others.

**Table 3.**
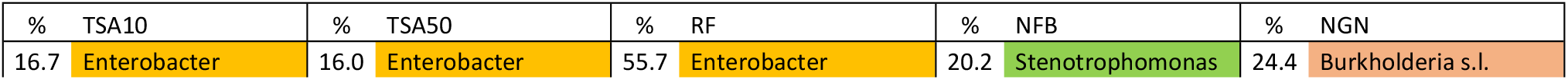
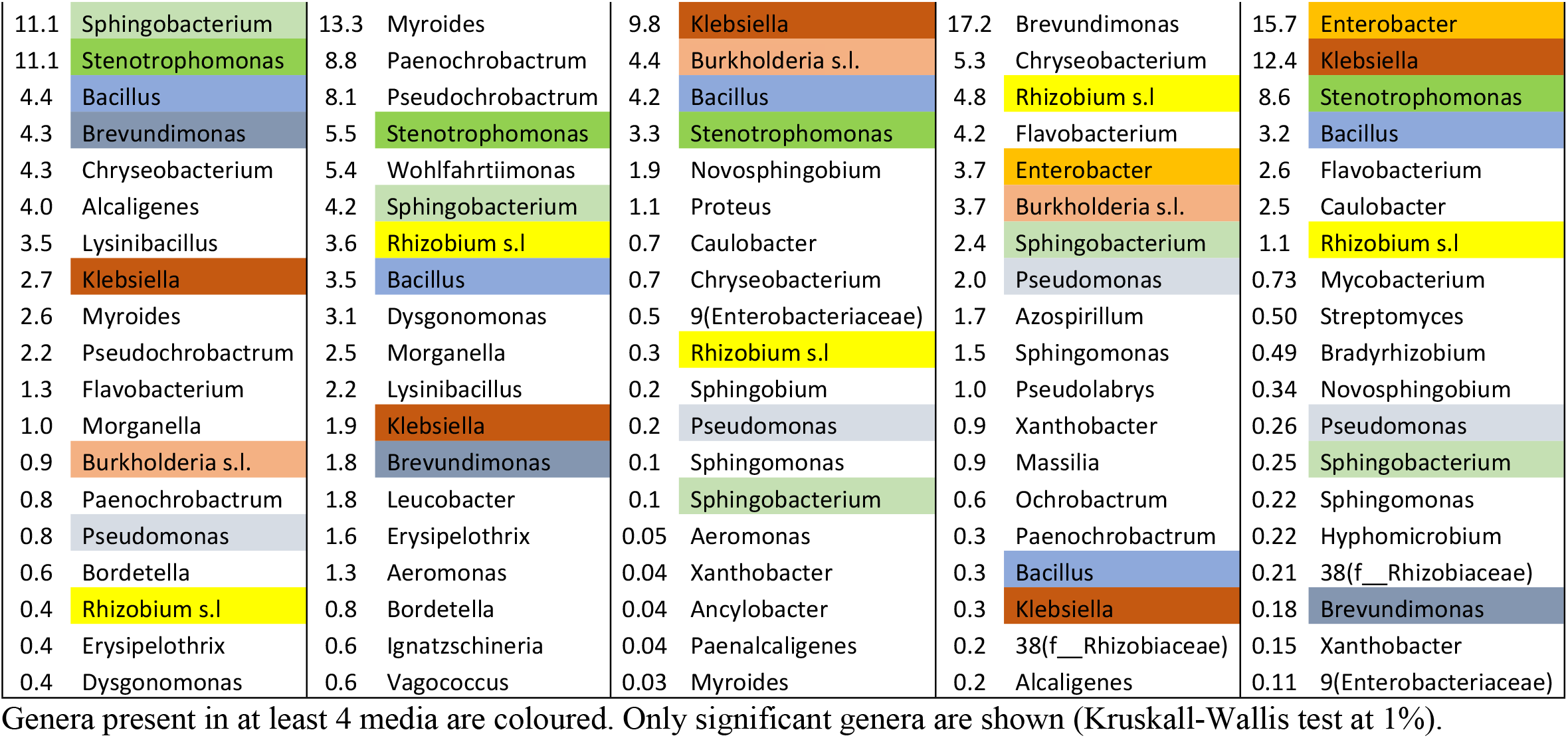
Distribution and mean relative abundance of top 20 genera detected in the CDA.

### Prediction of enriched functions in CIA compared to culture-based approach

We performed a functional prediction analysis using Picrust2 to infer metabolic capacities from our 16S amplicons ASV. Functions can be predicted in three classes: enzyme classification (EC), KEGG orthology (KO) and molecular pathways (PW). Data were normalised with relative abundance, and a Kruskal-Wallis test was performed across conditions (medium used for CDA and CIA). In order to evaluate the predictive ability of PiCrust2 algorithm on our dataset, we looked at the specific enzyme nitrogenase (EC. 1.18.6.1) prediction in the CDA libraries that included medium with (TSA, RF) or without nitrogen (NGN, NFb) (Figure 5A). As expected, we observed an enrichment of nitrogenase (p= 0.00492) in the nitrogen-free NFb and NGN media, with NGN medium exhibiting a much higher enrichment than NFb. The non-selective medium TSA and the plant-based medium (RF) did not enrich bacterial taxa with nitrogenase function (Figure 5A).

**Figure 5.**
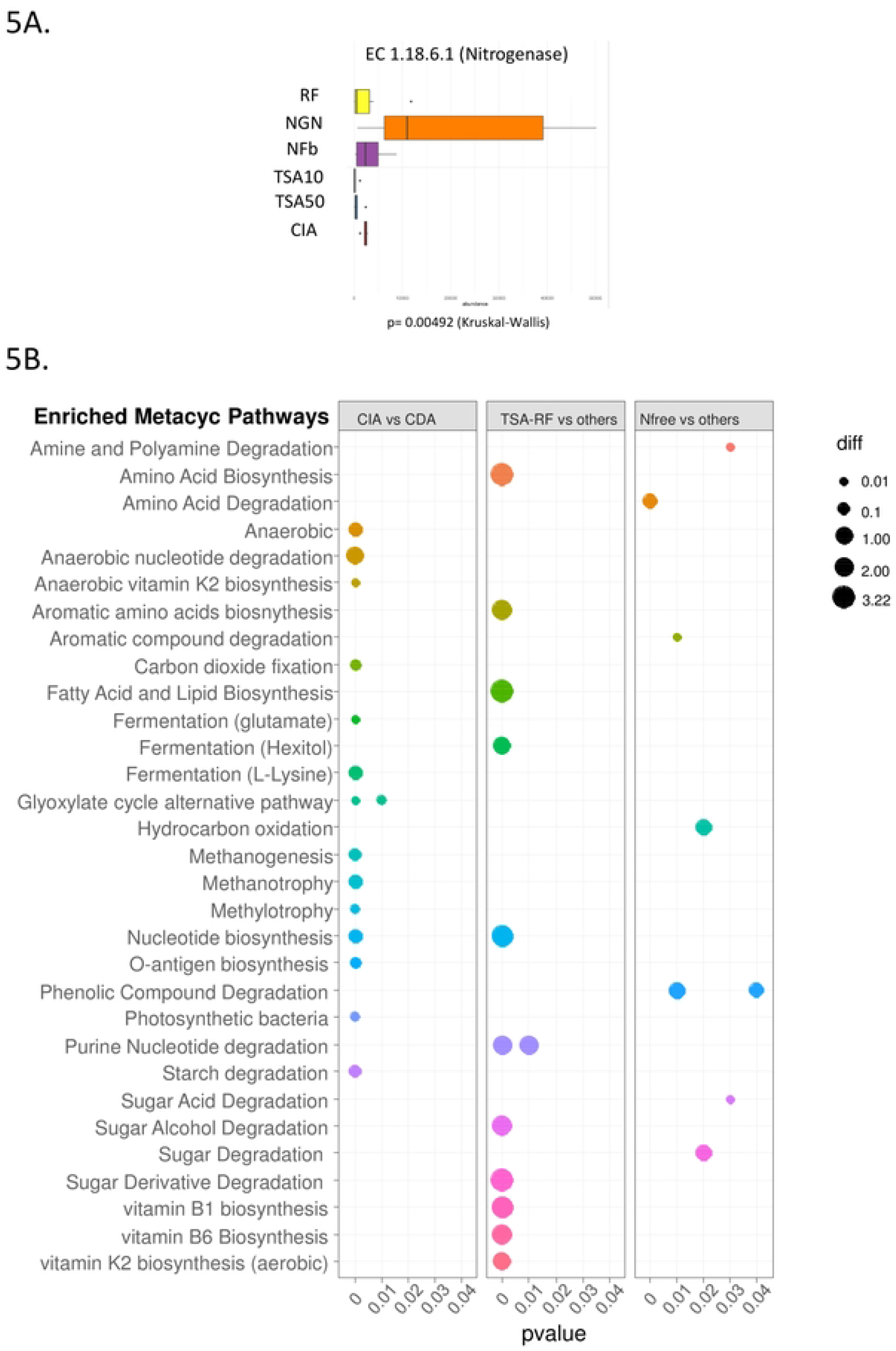
Nitrogenase enrichment prediction in 16S amplicon libraries (A) and dot-plot of predicted enriched Metacyc pathways (B) in cross comparisons CIA vs CDA, non-selective media (TSA, RF) versus others, and nitrogen-free media versus others.

We also aimed at predicting which functional pathways are specific of CIA compared to CDA, in order to help design conditions to capture the yet unculturable diversity. We thus analysed the metabolic pathways (based on PW/Metacyc categories) predicted as enriched in the CIA compared to CDA conditions, and represented the results in a dot-plot (Figure 5B). Among detected Metacyc pathways enriched in CIA, several functions linked to specific ecological niche abilities were detected: anaerobic/fermentation metabolism, carbon dioxide fixation, bacterial photosynthesis, methanotrophy and methylotrophy. Given our CDA culture conditions were aerobic and in the dark, such enrichment appears logical and give clues on culture conditions to capture more bacterial diversity. Enriched pathways in the TSA and RF media libraries (compared to others) could be linked to heterotrophy on rich media in aerobic conditions (sugar degradation, amino acid/lipid/nucleotide biosynthesis, vitamin biosynthesis). For the Nitrogen-free media (compared to others), several pathways were detected as phenolic compound/ polyamine/amino acid degradation and sugar degradation. It is to note that nitrogen fixation does not appear as itself in Metacyc pathways, it is embedded in “nitrogen metabolism” together with “nitrification” and “denitrification” capacities among others, so that no pattern from nitrogen fixation ability is possible, apart from the analysis on E.C. for the nitrogenase enzyme (Figure 5A).

## Discussion

In this study we aimed at quantifying the bacterial diversity bias observed when culturing rice-root associated bacteria on a range of culture media compared to real diversity, using Illumina sequencing on 16S-amplicon barcodes (variable region V3-V4). Our goal was to document precisely what is the bacterial taxa diversity that can be recovered from a set of culture media compared to real diversity, what predicted and enriched/depleted functions can be inferred from this diversity, and how to design new culture conditions to capture it. If several studies already compared rice microbiota culturable and real diversity (30–32), they often rely on comparisons between regular 16S Sanger sequencing on isolated bacteria compared to NGS sequences. Here by using the same sequencing methodology at high depth, we could compare diversity levels without sequencing/analytic bias. We also used two different analytic methods to infer operational taxonomic units, either ASV (based on exact sequence variants detection, (12) or OTU (based on clustering by swarming, (11)). As ASV analysis detected more diversity than OTU at different levels (even class and orders, Table 1), we preferred ASV for all subsequent diversity and functional predictions. As exact sequence variants strongly rely on algorithms for detection and correction of sequencing errors, we cannot exclude that some diversity obtained rely on imperfections of the algorithms. However, given that the obtained higher diversity concerns also higher taxonomic levels, the problem of such artificial diversity is unlikely as it would rely on a high number of mutations.

In this study the diversity obtained from the CDA culture media (TSA10, TSA50, RF, NGN, NFb) was lower compared to the microbiota (CIA). If we combine all diversity of the CDA, it represents 11.7% (ASV level), 29% (Genus level), 22.4% (Class level), 25.6% (Order level), 23.1% (Family level) of the diversity of the microbiota (CIA). In the literature, it is hard to find comparable studies to establish if our recovery rate is low or high, since this study is the first to our knowledge to have assess culturable recovery by amplicon barcoding and NGS sequencing. The review of Sarhan et al (16) detailed recent advances in culturomics methodologies, and establish to around 10 % the rate of recovery of conventional chemically-synthetic culture media, which is in the range of what we obtained at ASV level (though we obtained between 23 and 29% at upper levels of taxonomy). The study of (31) claim to recover up to 70% of bacterial species of the *Oryza sativa indica* and *japonica* rice microbiota, but they applied a cut-off on the frequency at 0.1%. If we apply the same cut-off on our dataset, we then detect 121 genera in the CIA of which 36 (29%) are present in the CDA.

From all media used in the CDA, we could recover a total of 142 bacterial genera, with each medium capturing 15 to 23 specific genera compared to each other (Figure 4C). The only exception is the “Rice Flour” medium, a plant-based media that captured here a low diversity of bacteria compared to others, probably due to its low complexity in composition. Plant-based media have been reviewed as good alternative to popular bacterial chemical media for increasing cultivability of plant-associated microbes (16), but they advise the use of homogenised roots, leaves, or exudates as complements to minimal or more complex media.

An unexpected feature observed from our data is the recovery of specific ASV from the CDA that were not detected in the CIA. This diversity represents 532 ASV, 1 class, 3 orders, 16 families and 70 genera (Table 1, Figure 4AB). One explanation for not detecting such diversity in the microbiota is the depths of sequencing, though the rarefaction curves did reach a plateau but at much higher alpha diversity for the CIA compared to CDA (Figure 2A). The sequencing depths obtained was in mean at 36120. If differences between bacterial ASV frequencies exceeds 10e^4^, then several genera may be undetected in the CIA approach while selected by specific culturable media. As we applied a filter on the number of reads at 10 (cumulated in all libraries), we also looked at lower filtering (>2) or unfiltered ASV data (Supplementary Table S5). Still we detected one specific class in CDA (undetected in CIA), which is *Erysipelotrichia* and is represented by one genus, *Erysipelothrix*, with 4 ASV recovered on TSA medium (at 10 and 50% concentration). A Blast of these ASV sequence revealed 100% sequence identity with the 16S rDNA of *Erysipelothrix inopinata*, a species which type strain was isolated from sterile-filtered vegetable broth (33). As our medium was sterilized by autoclaving, there is low probability that these 4 ASV were contaminants, and as the original species was in vegetable broth it seems it can be recovered from vegetable material. Another point to mention is that we are not the first to mention isolates from culturable approaches that were undetected in culture independent approaches (14,32), so it appears as frequent to recover bacterial strains from culturing approaches that are very low in frequency in microbiome data and even undetected. A way for better evaluating the diversity of rice microbiota would be to increase the sequencing depths to much higher degrees to have a better image of its whole diversity. In our study, differences in frequencies between ASV exceeding 10e^3^ (at genus level, Supplementary Table S4), would make them undetected in the CIA while observed in the culturomic approach. It appears critical since several studies have underlined the role of rare species (also called satellite taxa) in plant-microbe interactions and more broadly in key functions of the ecosystem (13,34). Another dimension requesting scientific efforts would be an increase in the representativeness of taxonomic diversity in databases as many ASV cannot be affiliated to taxonomic ranks due to missing descriptions of these taxa in taxonomic databases.

We also tried to predict functions and metabolic pathways that would be enriched when using different types of media, and did statistical tests to evaluate which functions are missing from our culturable approach. We found that many taxa with anaerobic metabolisms as methanogenesis (production of methane), methanotrophy (methane degradation) methylotrophy (one-carbon reduction) or photosynthetic capacities, were missing from our culturable approach (Figure 5). It is well known that the rice microbiota is different from others crops as it is often grown in flooded conditions, creating an oxic-anoxic interface between the rhizosphere/root system and the bulk soil (4,5). Our functional prediction approach thus underlined the presence of these bacteria adapted to anoxic conditions and probably strictly anaerobic in the CIA approach, and absent from the CDA. These predictions give clues on the specific conditions and composition of media to use to capture these yet unculturable functional groups of bacteria, and can be used to develop culturomics, a growing scientific field for microbiologist interested in synthetic microbiota and for biotechnological applications of plant-associated microorganisms.

## Acknowledgements

This work was performed thanks to the facilities of the “International Joint Laboratory LMI PathoBios: Observatory of plant pathogens in West Africa: biodiversity and biosafety” (www.pathobios.com; twitter.com/PathoBios). We are very grateful to Issouf Sanga, Laurentine Kabre, Sylvie S. Guissou, Alexandre P. Ouedraogo and Manaka Douanio for their contribution to this work. We also thank Gilles Bena, Pierre Czernic, Ludivine Guigard and Nicolas Busset for constructive discussions throughout this project. We thank the Institute of Research for Development (IRD) for awarding MS a PhD Research grant through its ARTS program.

## Supporting information Caption

Supplementary Table S1 : Dilutions and incubation time used for DNA extractions in the culturable approach.

Supplementary Table S2. 16S amplicons statistics in DADA2 pipeline.

Supplementary Table S3. Abondance of bacterial class detected in the rice microbiota (Wilcoxon test), and their occurrence in the culturable approach (Excel file).

Supplementary Table S4. Abondance of the 264 bacterial genera detected in the rice microbiota (Kruskal-Wallis test), and their occurrence in the culturable approach. (Excel file).

Supplementary Table S5. ASV count table with taxonomic ranks (sheet 1 :unfiltered, sheet 2 &3 filtered at 2 and 10 cumulated reads in all libraries, respecitvely) (Excel File).

Supplementary Figure S1. Richness (1A) and beta-diversity (1B) of root and rhizosphere 16S amplicon libraries.

